# Cytokine-Induced Transcriptional Changes in Human Neutrophils Reveal Immune Regulatory Plasticity

**DOI:** 10.1101/2025.02.20.639233

**Authors:** Huw B. Thomas, Steven W Edwards, Helen L Wright

## Abstract

Neutrophils are the major cellular constituent of blood leukocytes and play a central role in the inflammatory response, producing an array of destructive molecules and antimicrobial proteases that characterise the cells as front-line defenders of the innate immune system, crucial to host defence. It is now appreciated that neutrophils produce and respond to a variety of inflammatory signals and are able to regulate both the innate and adaptive immune responses. However, the mechanisms by which neutrophils respond to different inflammatory signals to regulate their own function and the functions of other immune cells are incompletely defined. In this study, we performed RNA sequencing with bioinformatics analysis of healthy human neutrophils exposed for 1h to a range of pro-inflammatory cytokines. GM-CSF and TNFα induced significant changes in expression in the most transcripts including activation of genes regulating apoptosis and genes encoding cytokines and chemokines that can drive the differentiation and activation of CD4 T cells. Stimulation of neutrophils with G-CSF, IFNα, IFNγ, IL-1β, or IL-8 resulted in expression of discrete gene sets and differential activation of signalling pathways including changes in cell adhesion and migration, immune receptor expression, apoptosis, and production of pro-inflammatory prostaglandins. This work defines the differential gene expression patterns in neutrophils exposed to different regulatory cytokines. This is important in both increasing our understanding of the role of neutrophils in driving innate and adaptive immune responses and, importantly, for deconvoluting the neutrophil gene expression signatures observed in inflammatory diseases.

## INTRODUCTION

Neutrophils are the most abundant leukocytes found in circulating blood and form the major cellular constituent of the innate immune system (1). They are indispensable for defence against invading bacterial and fungal pathogens due to their ability to phagocytose micro-organisms, release lytic enzymes from internal granules, produce reactive oxygen species (ROS) with antimicrobial potential and produce neutrophil extracellular traps (NETs) (2, 3). Their highly-conserved mechanisms of anti-microbial activity, coupled with a characteristic short lifespan, have historically defined neutrophils as a one-dimensional effector cell with little capacity to influence the more complex, adaptive arm of the immune system, predominantly regulated by T-cells and B-cells. However, in recent years, this view of neutrophils has been profoundly altered (4). Neutrophils are now known to produce and release numerous cytokines and chemokines plus angiogenic and fibrogenic factors (5-7). They have also been shown, following cytokine stimulation, to express MHC Class II molecules and present antigen to T-cells (8, 9). The cytokines tumour necrosis factor alpha (TNFα) and granulocyte/macrophage colony-stimulating factor (GM-CSF) are potent regulators of neutrophil gene expression, and both cytokines delay neutrophil apoptosis *in vitro* (10).

The perceived role of neutrophils in inflammatory disease has also been altered in recent years. Neutrophil dysregulation has been associated with the pathogenesis of a variety of cytokine-driven, chronic inflammatory diseases such as rheumatoid arthritis (RA), juvenile (and adult) systemic lupus erythematosus (SLE), chronic obstructive pulmonary disease (COPD), asthma, covid-19 and alcoholic hepatitis (2, 4, 11-18). Activated neutrophils are found at sites of inflammation such as the RA joint (19-21) as well as SLE skin lesions and regions in the kidney (22, 23). NET-producing neutrophils can be detected in RA blood smears (24). This current view of neutrophils places them central to the immune system with a significant capacity to regulate, influence and affect both the innate and adaptive response in health and disease. Despite a greater appreciation for neutrophil involvement in the immune response, relatively little work has focused on the underlying mechanisms of neutrophil activation and regulation in the context of inflammation, instead focusing more on the traditionally associated mechanisms of functions such as chemotaxis, phagocytosis and apoptosis. Recent transcriptomic studies on inflammatory neutrophils have identified a key role for interferon-induced genes in the auto-immune diseases RA and SLE (25, 26), and several metabolomics studies have also shed insight into the key roles of glycolysis, oxidative phosphorylation and amino acid recycling in inflammatory neutrophils (27, 28). Single-cell studies have revealed the heterogeneity of neutrophil subsets in homeostasis and inflammation, and sub-sets of neutrophils including low-density granulocytes (LDGs) can be distinguished by their gene expression profiles (29-31). Epigenetic mechanisms, such as DNA methylation and histone modifications, play crucial roles in regulating gene expression in neutrophils (32), and chromatin remodelling under conditions of inflammation can induce expression of interleukin-6 (IL-6), IL-12B and IL-23A by human neutrophils (33, 34).

The aim of this study was to measure the rapid changes in gene expression in healthy human neutrophils that are induced by inflammatory cytokines in order to determine the role of neutrophils in immune regulation during inflammation. We selected a range of colony-stimulating factors (CSF), interferons (IFN), interleukins (IL) and TNFα for investigation, and show that these individual cytokines induce expression of distinct genes regulating a range of signalling pathways, including expression of their own respective negative signalling regulators. We also show that neutrophils can express genes for a range of cytokines, chemokines and immune receptors involved in driving both innate and adaptive immune responses.

## METHODS

### Ethics and patients

This study was approved by the University of Liverpool Central University Research Ethics Committee. All participants gave written, informed consent in accordance with the declaration of Helsinki. Healthy controls were recruited from staff at the University of Liverpool. All participants were over the age of 18 years and free of infection. All samples were collected between 09.00-10.00 to avoid any differences in gene expression that may be caused by circadian rhythms (35-37),

### Neutrophil isolation and culture

Neutrophils were isolated from heparinised peripheral blood using Polymorphprep (Axis Shield) as previously described (38). Contaminating erythrocytes were lysed using ammonium chloride lysis buffer. Neutrophils were resuspended in RPMI 1640 media (Life Technologies) containing L-glutamine (2mM) and HEPES (25mM) at a concentration of 5×10^6^/mL unless otherwise stated. Neutrophil purity was routinely >97% and viability >98%. Neutrophils were incubated with inflammatory cytokines (Merck) for 1h prior to preparation of RNA lysates, as follows: granulocyte colony-stimulating factor (G-CSF), interferon alpha (IFNα), interferon gamma (IFNγ), interleukin-1 beta (IL1β), TNFα (all 10ng/mL), GM-CSF (5ng/mL), interleukin-8 (IL-8, 100ng/mL).

### RNA isolation

RNA was isolated from 5-10×10^6^ neutrophils using an optimised Trizol-chloroform protocol (Life Technologies) (10, 38), precipitated by isopropanol and cleaned using the Qiagen RNeasy (mRNA) kit including a DNase digestion step. RNA was snap-frozen in liquid nitrogen and stored at -80°C. Total RNA concentration and integrity was assessed using the Agilent 2100 Bioanalyser RNA Nano chip. RNA integrity was routinely ≥ 7.0.

### mRNA sequencing

Total RNA was enriched for mRNA using poly-A selection. Fifty base pair single-end read libraries were sequenced on the Illumina HiSeq 2000 platform. Reads were mapped to the human genome (hg38) using Tophat2 (v2.04) (39). Read counts were generated using featureCounts (Rsubread package v2.0.1) (40) for R (v4.0.2) (41). Statistical analysis of gene counts was carried out using DESeq2 (v1.44) (42) and limma (v3.60.0) (43) in R.

### Bioinformatics and statistical analysis

Gene ontology enrichment analysis of genes with an adjusted p-value ≤0.05 between each cytokine vs untreated was carried out using the R package clusterProfiler against the genome wide annotation for human (org.Hs.eg.db) (44, 45). Lists of gene identities within ontologies were downloaded from the Gene Ontology Resource (https://geneontology.org). Functional enrichment analysis of genes with an adjusted p-value ≤0.05 and fold-change ≥2 or ≤-2 between each cytokine vs untreated, was complemented with Ingenuity Pathway Analysis (https://digitalinsights.qiagen.com/IPA) (46) canonical pathway and upstream regulator analysis, using the Ingenuity Knowledge Base as background. A full list of R packages used in our analyses can be found in Supplementary Data.

### Flow cytometry

Following incubation of 10^6^/mL neutrophils with cytokines at the concentrations listed above for 22h at 37°C in 5% CO_2_, 10^5^ cells were incubated in 100 μL HBSS with 10 μg/mL of FITC-conjugated annexin V (Merck), in the dark. After 15 min, 1 μg/mL propidium iodide (PI, Merck) was added and samples were measured immediately on a Guava Easycyte flow cytometer. A minimum of 5000 gated events were collected per sample.

### Western blotting

Following incubation of 5×10^6^/mL neutrophils with cytokines at the concentrations listed above for 15min at 37°C in 5% CO_2_, cells were centrifuged at 1000 g for 3 min. Cell pellets were rapidly lysed in boiling Laemmli buffer for 5 min. Neutrophil lysates (10µL) were separated by SDS-PAGE (10-15% depending on protein size) before transfer to PVDF membranes, which were blocked for 1h using 5% non-fat dry milk. Primary antibodies (in 5% BSA) used to detect phospho-proteins were: p-STAT1, p-STAT3, p-NFκB (p65), p-AKT, p-ERK, p-p38 MAPK (1:1,000 dilution, all from Cell Signalling), and actin (1:10,000 dilution, Merck). Secondary antibodies (1:10,000) were from Merck. Bound antibodies were detected using enhanced chemiluminescence (ECL) reagents (Merck) and careful exposure of the membrane to hyper-film in a dark-room.

## RESULTS

### In vitro cytokine treatments induce distinct gene expressions in human neutrophils

Principal Component Analysis (PCA) of cytokine-treated neutrophil RNA-Seq datasets identified clustering of samples based on cytokine stimulus (Figure 1A). GM-CSF induced the highest number of significant differently expressed (DE) genes compared to untreated neutrophils, with 3,728 genes having an adj. p-value <0.05. TNFα treatment resulted in 2,134 DE genes, followed by IFNγ with 1,331 and G-CSF with 1,300. IL-8 significantly altered 1,229 genes, IFNα 210 and IL-1β 46 genes. These DE genes are summarised in a heatmap (Figure 1B). Comparison of the DE genes using an upset plot (Figure 1C) identified overlapping DE genes between different cytokine treatments. GM-CSF and TNFα had the most common number of DE genes at 553, with 317 DE genes being shared between GM-CSF and IL-8. The upset plot also provides details of the number of DE genes uniquely altered by each cytokine treatment compared to untreated neutrophils.

**Figure 1.**
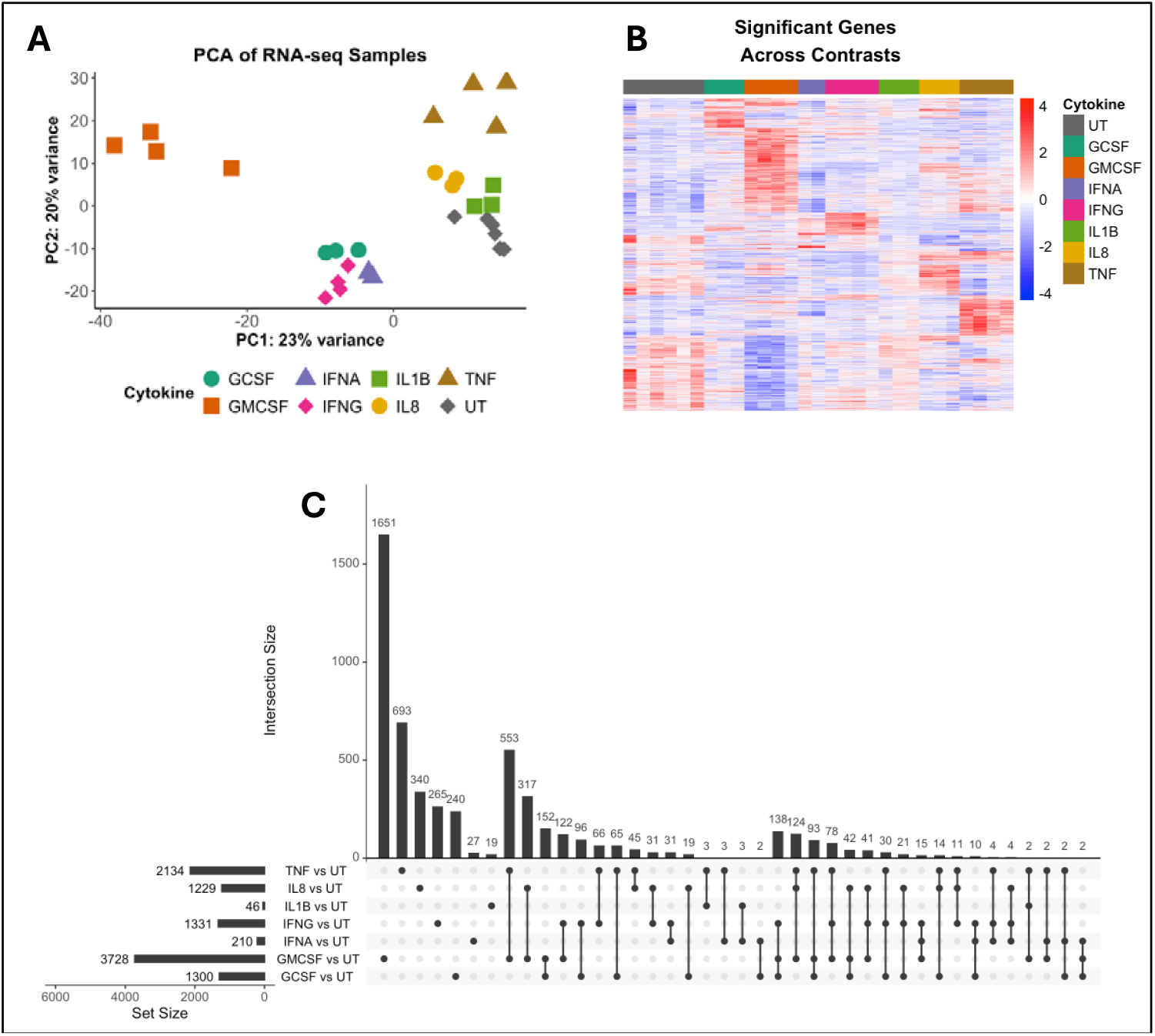
Transcriptome analysis of cytokine-treated human neutrophils. (A) Principal Component Analysis of whole transcriptomes from neutrophils treated with G-CSF, GM-CSF, IFNα, IFNγ, IL-1β, IL-8 or TNFα or left untreated (UT) for 1h. (B) Heatmap of differentially expressed (DE) genes across all samples (adj. p-value<0.05). (C) Comparison of DE genes between cytokine treatments (adj. p-value<0.05). Set size indicates total number of DE genes, and intersection size indicates number of DE genes common to the treatments connected by black dots.

### Gene ontology analysis of cytokine-treated neutrophils reveals altered effector functions

Volcano plots for each cytokine-treated comparison against untreated neutrophils is shown in Figure 2A. Gene Ontology enrichment analysis was performed using clusterProfiler and is summarised in Figure 2B. G-CSF and GM-CSF increased expression of a number of genes which can regulate T cell activation predicting enrichment of T cell differentiation, CD4 T cell activation, Th1 immune functions and regulation of cytokine production. Expression levels for these genes are shown in a heatmap (Supplementary Figure 1A). IFNα DE genes were strongly enriched for GO Terms relating to response to viral infections and regulation of innate immune responses (Supplementary Figure 1B), whereas IFNγ DE genes were enriched for regulation of the innate immune response and the humoral immune response. IL-1β upregulated genes involved in neutrophil chemotaxis and migrtion as well as the cellular response to a biotic stimulus (Supplementary Figure 1C). IL-8 responses included prostaglandin transport and secretion (Supplementary Figure 1D). TNFα DE genes were enriched in the most GO terms, including regulation of cytokine production, cell adhesion, regulation of CD4 T cell activation, neutrophil migration, as well as GO terms indicating response to TNF and canonical NF-κB signal transduction (Supplementary Figure 2A,B).

**Figure 2.**
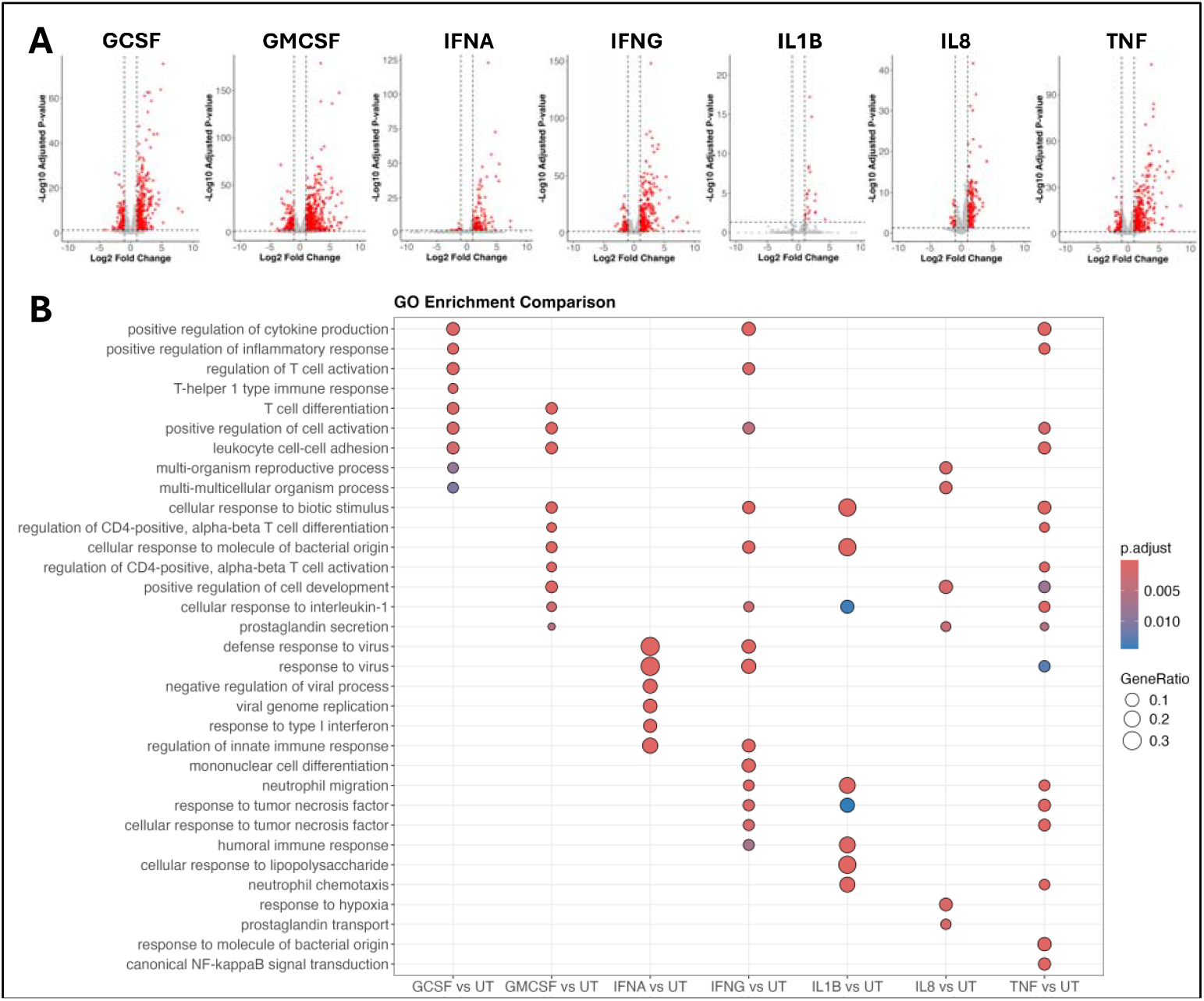
Gene Ontology enrichment analysis of cytokine-treated human neutrophils. (A) Volcano plots showing genes significantly up- and down-regulated (red, adj. p-value < 0.05) in neutrophils treated with G-CSF, GM-CSF, IFNα, IFNγ, IL-1β, IL-8 or TNFα for 1h compared to untreated (UT) neutrophils. (B) Summary of Gene Ontology (Biological Process) enrichment analysis of cytokine-treated neutrophils.

### Upregulation of pro-inflammatory and anti-apoptotic genes in cytokine-treated neutrophils

We observed that expression of genes for cytokines and chemokines was significantly different between cytokine-treated neutrophils. GM-CSF and TNFα up-regulated the most genes for cytokines (including IL1A and IL1B, Figure 3A) and chemokines (including CXCL1, CXCL2 and CXCL8). TNFα treatment upregulated its own expression. IFNα and IFNγ both upregulated expression of CXCL9, CXCL10 and TNFSF10, whereas G-CSF and IFNα increased expression of TNFSF13B.

**Figure 3.**
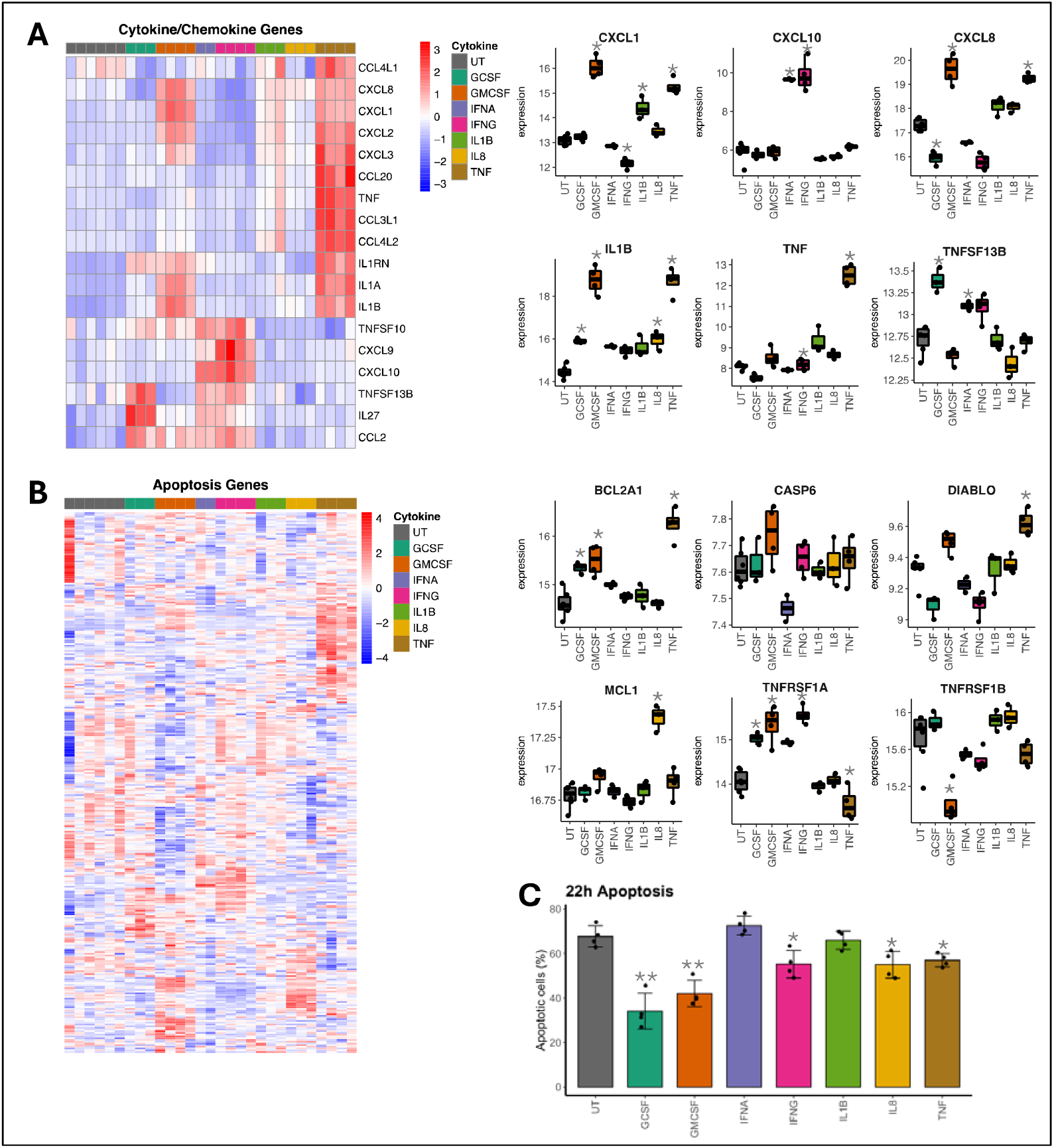
Analysis of cytokine production and apoptosis genes. (A) Heatmap showing expression of genes encoding cytokines and chemokines in neutrophils treated with G-CSF, GM-CSF, IFNα, IFNγ, IL-1β, IL-8 or TNFα for 1h compared to untreated (UT) neutrophils. Box plots of selected genes are shown (* adj. p-value<0.05). (B) Heatmap showing expression of genes which regulate apoptosis in neutrophils treated with G-CSF, GM-CSF, IFNα, IFNγ, IL-1β, IL-8 or TNFα for 1h compared to untreated (UT) neutrophils. Box plots of selected genes are shown (* adj. p-value<0.05). (C) Levels of neutrophil apoptosis measured by annexinV/PI flow cytometry in healthy neutrophils after 22h incubation with cytokines (* p<0.05, ** p<0.01).

The expression of apoptotic genes was also differentially regulated by cytokine-treatment. The anti-apoptotic gene BCL2A1 was upregulated by G-CSF, GM-CSF and TNFα, and anti-apoptotic MCL1 was upregulated by IL-8 (Figure 3B). Caspase 6 and DIABLO were up-regulated by GM-CSF and TNF respectively. There was differential regulation of the two TNF receptors; TNFRSF1A was up-regulated by G-CSF, GM-CSF and IFNγ and this was associated with a significant down-regulation of TNFRSF1B. Changes in expression of apoptotic genes was closely mirrored by measurement of apoptotic neutrophils in culture after 22h cytokine treatment (Figure 3C). G-CSF, GM-CSF and IFNγ were the most anti-apoptotic cytokine treatments, with IL-8 and TNF being moderately anti-apoptotic. IFNα and IL-1β had no significant effect on neutrophil apoptosis.

### Signalling pathway analysis of cytokine-treated neutrophil transcriptomes

We used Ingenuity Pathway Analysis (IPA) to predict the canonical signalling pathways up- and down-regulated in cytokine-treated neutrophils. A comparison analysis of the most significantly altered canonical pathways is shown in Figure 4A. A number of these pathways were common to several cytokine treatments, for example Class A/1 rhodopsin-like receptor signalling, FXR/RXR activation, G alpha (ii) and G-protein coupled signalling, pathogen induced cytokine storm and S100 family signalling (adj. p-value <0.05). TREM1 and PPAR signalling was uniquely up-regulated in TNFα-treated neutrophils. p38 MAPK and HMGB1 signalling was upregulated in GM-CSF and TNFα treated neutrophils, and unsurprisingly interferon alpha/beta signalling was up-regulated in IFNα and IFNγ signalling. The peroxisome proliferator-activated receptor (PPAR) signalling pathway was down-regulated by all cytokine treatments, whereas integrin cell-surface interactions was down-regulated by all cytokines except TNFα.

**Figure 4.**
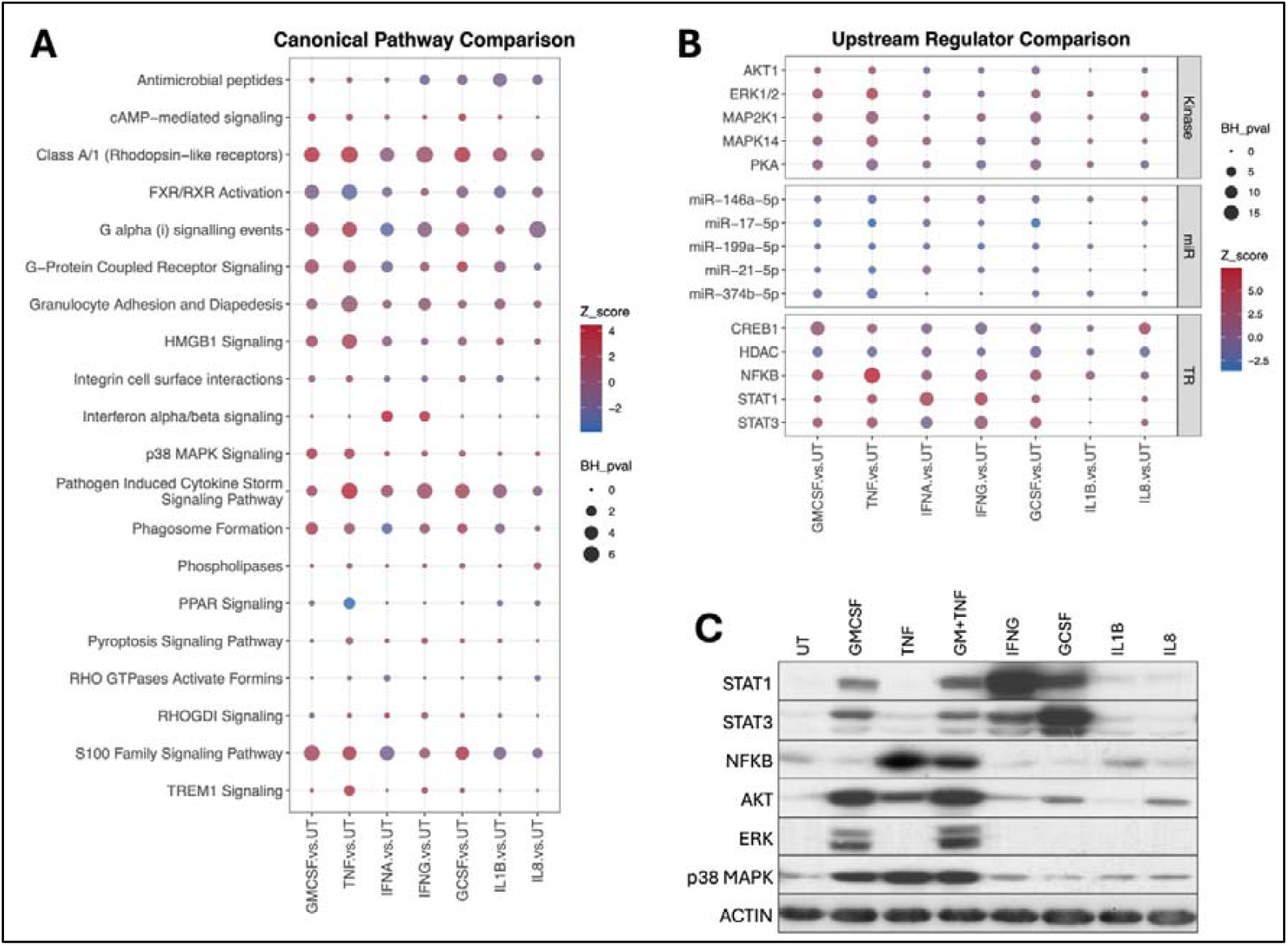
Ingenuity pathway analysis of neutrophil transcriptomes. (A) Summary of canonical signalling pathways up- and down-regulated in neutrophils treated with G-CSF, GM-CSF, IFNα, IFNγ, IL-1β, IL-8 or TNFα for 1h compared to untreated (UT) neutrophils. (B) Summary of upstream regulators predicted to be activating neutrophils neutrophils treated with G-CSF, GM-CSF, IFNα, IFNγ, IL-1β, IL-8 or TNFα for 1h compared to untreated (UT) neutrophils (miR, microRNA; TR, transcription regulator). (C) Western blot for phosphorylated signalling proteins in cytokine-treated neutrophils after 15 min. GM-CSF+TNFα shown as positive control lane.

IPA up-stream regulator analysis was used to predict which kinases, microRNAs (miR) and transcription regulators (TR) were regulating the cytokine-treated neutrophil transcriptomes (Figure 4B). This identified AKT1, ERK1/2, MAP2K1, MAPK14 and PKA as key kinases activated by a number of cytokines in human neutrophils. Also predicted was strong activation of NF-κB in TNFα-treated neutrophils, STAT1 and STAT3 in interferon-treated and CSF-treated neutrophils, and CREB1 in G-CSF, GM-CSF, IL-1β and IL-8 treated neutrophils. HDAC was predicted to be down-regulated by all cytokine treatments. Differential regulation of microRNAs was also predicted. We validated the activation of a number of these transcription regulators by western blotting for phospho-proteins STAT1, STAT2, NF-κB (p65), AKT, ERK(1/2) and p38 MAPK (MAPK14) in neutrophils treated with cytokines for 15 min (Figure 4C).

### Altered expression of immune-receptors in cytokine-treated neutrophils

We identified significant differences in the expression of genes for immune-receptors, including cytokine and chemokine receptors, complement receptors, Fc receptors and genes encoding MHC Class I and II (Figure 5A). FcγR1A was uniquely upregulated by IFNγ. FcγR2B was up-regulated by G-CSF and GM-CSF, and FcγR3b was down-regulated by GM-CSF, IL-8 and TNFα. The chemokine receptors CXCR1 and CXCR2, through which CXC chemokines including IL-8 signal, were up-regulated by G-CSF and down-regulated by TNFα. CXCR4, which is involved in the homing of aged neutrophils back to the bone marrow for apoptosis, was down-regulated by all cytokines except IL-1β, supporting the results of the apoptosis assay (Figure 3C). Expression of the IFNα receptor gene IFNAR1 was increased by G-CSF, IFNγ and TNFα, and expression of the IFNγ receptor IFNGR1 was down-regulated by IFNγ, IL-1β, IL-8 and TNFα. The MHC Class II gene HLA-DRA was up-regulated by GM-CSF and TNFα.

**Figure 5.**
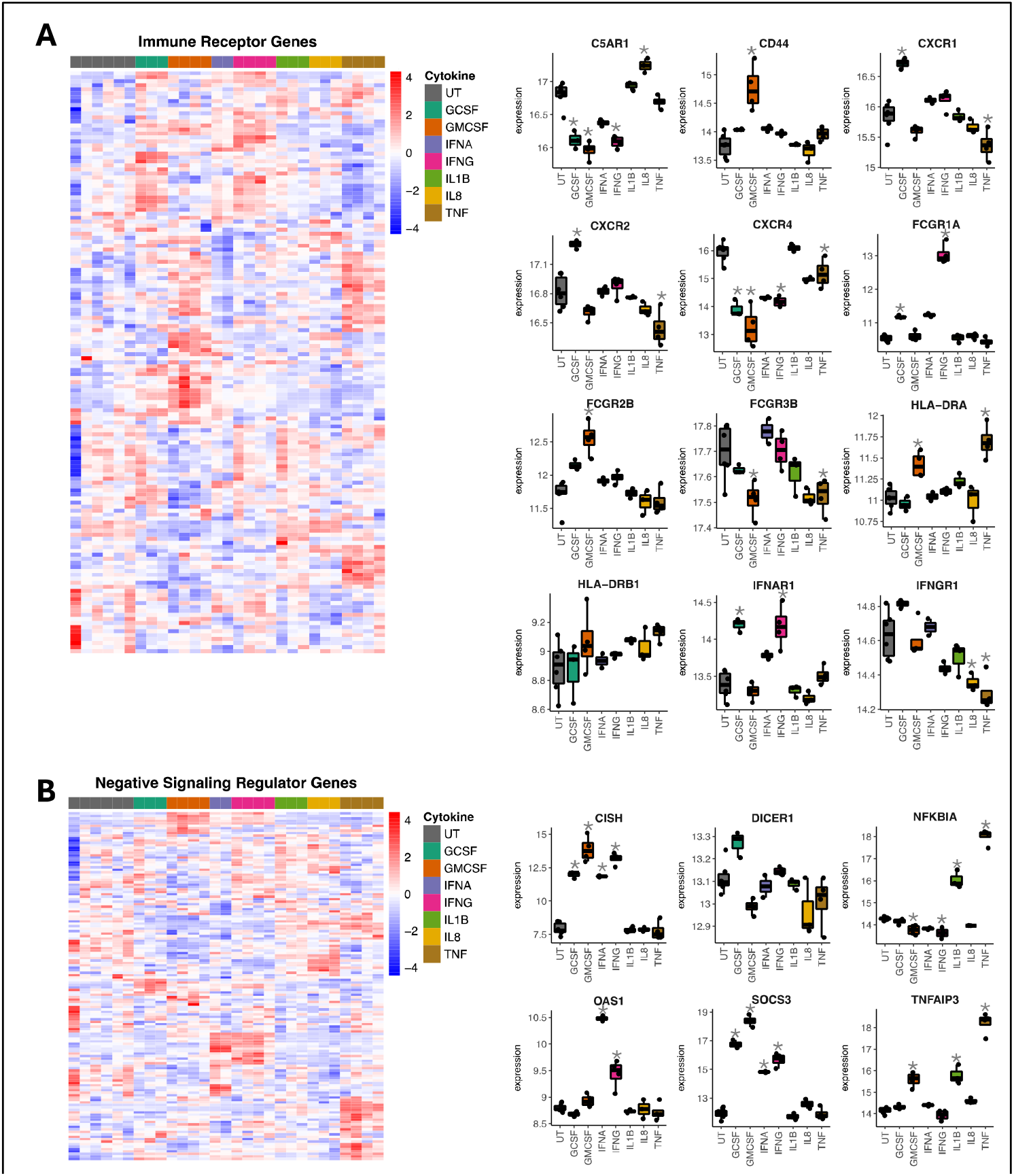
Analysis of immune receptor and negative regulators of cytokine signalling genes. Heatmaps showing expression of genes encoding (A) immune receptors and (B) negatively regulate cytokine signalling in neutrophils treated with G-CSF, GM-CSF, IFNα, IFNγ, IL-1β, IL-8 or TNFα for 1h compared to untreated (UT) neutrophils. Box plots of selected genes are shown (* adj. p-value<0.05).

### Negative regulation of cytokine signalling induced in cytokine-treated neutrophils

Whilst the cytokines used in this study induced activation of neutrophils, delay of apoptosis and expression of pro-inflammatory genes, they also induced activation of expression of negative regulators of cytokine signalling, indicating a negative feedback loop in human neutrophils. TNFα and IL-1β induced high expression of inhibitors of NF-κB signalling including NFKBIA and TNFAIP3. Cytokines which signal through the JAK/STAT pathway, including G-CSF, GM-CSF, IFNα and IFNγ, induced high expression of CISH and SOCS3. IFNα and IFNγ induced expression of OAS. In all cases, these negative regulators of cytokine-induced signalling pathways were amongst the most significant and up-regulated genes for each cytokine treatment. A key gene regulating microRNA processing, DICER1, was upregulated by G-CSF and down-regulated by GM-CSF and IL-8.

## DISCUSSION

The aim of this research was to measure the global changes in gene expression profiles of human neutrophils stimulated with a variety of inflammatory cytokines for 1h in order to define the role that these cells play in immune cell regulation and control of inflammation. The cytokines chosen for investigation have previously been shown to alter neutrophil functional responses e.g. apoptosis and/or priming (10, 47-50), and are often found at high concentrations in inflammatory biofluids such as RA synovial fluid (51). Thus, they represent both the trigger for activated neutrophil gene expression signatures *in vivo* and are potentially a target for the development of antibody-based or cell signalling inhibitor therapies in the future. In this study, neutrophils were stimulated with a range of cytokines/chemokines commonly associated with inflammation and found both systemically and at sites of inflammation. We measured differential regulation of thousands of genes in human neutrophils, with both cytokine-specific gene expression signatures and commonly-upregulated gene expression signatures identified. Our Gene Ontology enrichment analyses revealed a crucial role for inflammatory neutrophils in driving activation of immune responses, in particular in regulating the differentiation and activation of T cells. G-CSF, GM-CSF and TNFα all induced strong expression of neutrophil products involved in T cell activation, in particular CD4 Th1 cell activation. These Th1 cells, which produce high levels of TNFα and IFNγ, are particularly implicated in the pathology of RA, SLE and Crohn’s disease (52-55) where neutrophils are also shown to contribute to pathology, highlighting the interdependency and co-regulation of the innate and adaptive immune systems during cytokine-driven inflammation. Incubation of neutrophils with IFNα and IFNγ resulted in gene expression changes that mimicked the cellular defence response to viral infection, whereas IL-1β and TNFα activated genes that are typically involved in the cellular response to bacterial infection. IL-8 strongly increased expression of genes involved in the synthesis of pro-inflammatory prostaglandins. These included genes for prostaglandins PTGS2 (cyclooxygenase-2), PTGES (prostaglandin E synthase), as well as EDN1 (endothelin 1) which has been shown to induce the production of MMP9, TNFα, VEGF and IFNγ by neutrophils (56), and prostaglandin E2 by monocytes (57). This underlines the dynamic response of neutrophils to the inflammatory environment, and their rapid synthesis of pro-inflammatory molecules to help drive the specific innate and adaptive immune response required to clear an infectious threat.

Analysis of the effects of cytokine stimulation on neutrophil apoptosis revealed significant differences in the way these different molecules delay neutrophil life-span. We previously reported that both GM-CSF and TNFα delay neutrophil apoptosis, and that this is regulated through differential activation of two independent signalling pathways, JAK/STAT for GM-CSF and NF-κB signalling for TNFα treatment (10). It has also been shown by ourselves and others that human neutrophils do not express the anti-apoptotic gene BCL2, instead relying on MCL1 and BCL2A1 (Bfl1) for the regulation of apoptosis (50, 58-60). In this study we found that G-CSF, as well as GM-CSF and TNFα, had the strongest anti-apoptotic effect which was associated with increased expression of BCL2A1. IL-1β and IFNα had no effect on neutrophil apoptosis. Since neutrophils have a much shorter lifespan than other leukocytes, regulation of neutrophil apoptosis is an important biological process which may be critically-linked to activation of gene expression, for example, following migration into tissues. Cytokine activation of neutrophils at sites of inflammation leads to the enhanced expression of other chemokines and cytokines, as well as other key molecules such as adhesion molecules (19). Therefore, an extended neutrophil lifespan goes hand-in-hand with extended proinflammatory functions as part of the inflammatory response. These newly-expressed genes may themselves be directly involved in apoptosis delay for example via autocrine signalling, although it must be pointed out that regulation of apoptosis is also achieved by phosphorylation and stabilisation of anti-apoptotic proteins such as Mcl-1, without the requirement of *de novo* gene expression (59, 60).

A major finding of this study was that we identified significant increases in expression of genes encoding for a range of inflammatory cytokines and chemokines in cytokine-treated neutrophils. Importantly, neutrophils responded to exposure to different cytokines by increased expression of diverse sets of cytokines/chemokines that had differential effects on immune regulation. The importance of neutrophil-derived molecules during inflammation has long been overlooked in favour of cells of the adaptive immune response (largely B-cells, and T-cells). Indeed, much of the current knowledge on neutrophil-derived products is based on studies from non-human species (often mice), such that their production by human neutrophils remains without consensus in the scientific community (33, 34, 49, 61, 62). Our analysis of expressed cytokine/chemokine genes revealed both similarities and differences in the expression profiles of neutrophils stimulated with different cytokines. Perhaps unsurprisingly, conditions which had exhibited greatest ability to delay apoptosis also showed a similar pattern of cytokine expression, especially GM-CSF and TNFα stimulation. TNFα treatment led to a more than 10-fold increase in expression of TNFα and CXCL2 mRNA. Whereas GM-CSF treatment resulted in a >10-fold expression of CXCL1 and OSM, both treatments upregulated expression of IL1A, IL1B, and IL1RN. However, several chemokine genes showed decreased expression following IFNγ treatment compared to untreated control, for instance, CXCL1, CCL3, CCL4 and CXC12. Interferons -α and -γ uniquely upregulated expression of CXCL10, an important cytokine in the chemoattraction of adaptive immune cells (47). Thus, cytokine/chemokine production appears to be differentially regulated by these two inflammatory cytokines. These findings have important implications for inflammatory disease where high levels of inflammatory cytokines and neutrophils are common, such as TNFα in RA (51), and IFNα in SLE (63) and covid-19 (15, 17).

Among the genes with highest expression following treatment with any of the cytokines studied here were genes associated with suppression or inhibition of the signalling pathway that had been activated. For example, TNFα treatment induced expression of NFKBIA, NFKBIE and TNFAIP3 which are inhibitors of the NF-κB signalling pathway (64). Similarly, G-CSF, GM-CSF, IFNα and IFNγ increased expression of genes associated with suppression of JAK/STAT signalling (CISH and SOCS3) (65). This mechanism thus provides a mechanism to control and resolve cytokine-induced neutrophil signalling by up-regulation of a negative feedback loop which subsequently de-activates the neutrophils.

The importance of pro-inflammatory cytokines in systemic, autoimmune diseases is highlighted by the success of the use of anti-cytokine (or cytokine-receptor) drug therapy. Therapeutics such as Anakinra (IL-1R antagonist), Tocilizumab (anti-IL-6 receptor), Secukinumab (anti-IL17A) and Belimumab (anti-B-cell activating factor (BAFF)) are routinely used in a variety of inflammatory diseases such as gout, SLE, psoriasis, Crohn’s disease and RA. Newer, small molecule signalling inhibitors have been introduced more recently, the most successful of which are JAK inhibitors such as baricitinib (66). However, the most successful target for treatment of immune-mediated inflammatory disease is TNFα. Several drugs, such as Adalimumab, Cerolizumab-pegol and Etanercept, as well as biosimilars, are considered the front-line treatment for conditions such as RA, Ankylosing spondylitis and Crohn’s disease. However, an important feature of these drugs is the varying degree to which patients respond. For example, it is estimated that approximately 30% of patients with RA will not achieve adequate disease control with their first TNF inhibitor (67, 68). These patients will often have to switch therapies a number of times to alternative anti-TNF drugs, or to drugs targeting different proteins, such as Rituximab (anti CD20+ B-cell) or Abatacept (anti-CTLA4+ T-cell), before disease remission is achieved and maintained. This highlights the heterogeneity that exists in inflammatory diseases such as RA and suggests that different cytokines may be responsible for driving inflammation in different patients. Whilst treating inflammatory diseases using a single anti-cytokine drug is of merit, a comprehensive understanding of the molecular changes induced by inflammatory cytokines in health and disease, and how this regulation differs between cytokines and individuals is important to understand not just immune regulation, but could also lead to a more rationale-based approach to drug treatment.

We recognise some limitations in our study. Firstly, we only measured the change in gene expression induced by cytokines at a 1h timepoint. This has previously been shown to be an optimal timepoint to measure immediate changes in neutrophil gene expression in response to cytokines, likely as a result of activation of pre-existing transcription factors (50). We chose this timepoint as we wanted to measure early gene expression in response to a single cytokine. Later timepoints may be confounded by secondary responses to secreted products e.g. genes induced by autocrine production of, and response to, IL-8 or TNFα. Both these cytokines have been proven to be synthesised and secreted by activated neutrophils, inducing autocrine signalling and chromatin remodelling leading to the expression of IL-6 [33]. We also wanted to avoid later timepoints as these may have been affected by the presence of apoptotic neutrophils. Secondly, we were not able to perform all cytokine treatments on neutrophils from every donor, and this is reflected by having 6 untreated samples and 2-4 replicates in cytokine-treated sample. We additionally did not investigate the effect of combinations of cytokines (e.g. TNFα+IL-8) as part of this study, nor did we measure the response of neutrophils to pathogen associated molecular patterns such as lipopolysaccharide, or bacterial RNA or DNA. Importantly, we performed all our experiments using healthy donors and included both male and female neutrophil donors in the study.

In summary, our analysis of neutrophils under different conditions of simulated inflammation by different cytokine stimulation using RNA-Seq has revealed that neutrophils express discrete sets of genes in response to different stimuli. Analysis of the genes expressed reveals that several signalling pathways and transcription factors are differentially activated, which is confirmed by western blot analysis. Among the genes expressed are genes associated with cytokine-and chemokine-signalling which show differential expression among treatment conditions, and hence the ability of activated neutrophils to regulate different innate and adaptive immune responses. The data presented here reveal the plasticity of neutrophils under conditions of inflammation, and highlight the importance of proximal signals on the developing phenotype of a neutrophil during different forms of activation.

## ACKNOWLEDGEMENTS

We would like to thank all study participants for the generous donation of the blood samples used in this research. This work was undertaken on Barkla, part of the High Performance Computing facilities at the University of Liverpool, UK.

