## Supplementary Data for "Cytokine-Induced Transcriptional Changes in Human Neutrophils Reveal Immune Regulatory Plasticity"

List of R packages used in this publication:

Carlson M. (2025) Genome wide annotation for Human, primarily based on mapping using Entrez Gene identifiers. DOI: [10.18129/B9.bioc.org.Hs.eg.db](https://doi.org/doi:10.18129/B9.bioc.org.Hs.eg.db)

Conway JR, Lex A, Gehlenborg N. (2017). UpSetR: an R package for the visualization of intersecting sets and their properties. Bioinformatics. 33(18):2938-2940. doi: 10.1093/bioinformatics/btx364. PMID: 28645171; PMCID: PMC5870712.

Kolde R (2018). pheatmap: Pretty Heatmaps. R package version 1.0.12, <https://github.com/raivokolde/pheatmap>

Love MI, Huber W, Anders S (2014). Moderated estimation of fold change and dispersion for RNA-seq data with DESeq2. Genome Biology, 15, 550. [doi:10.1186/s13059-014-0550-8](https://doi.org/10.1186/s13059-014-0550-8).

Ritchie ME, Phipson B, Wu D, Hu Y, Law CW, Shi W, Smyth GK (2015). limma powers differential expression analyses for RNA-sequencing and microarray studies. Nucleic Acids Research, 43(7), e47. [doi:10.1093/nar/gkv007](https://doi.org/10.1093/nar/gkv007).

Wickham H (2016). ggplot2: Elegant Graphics for Data Analysis. Springer-Verlag New York. ISBN 978-3-319-24277-4, <https://ggplot2.tidyverse.org>

Wu T, Hu E, Xu S, Chen M, Guo P, Dai Z, Feng T, Zhou L, Tang W, Zhan L, Fu x, Liu S, Bo X, Yu G (2021). clusterProfiler 4.0: A universal enrichment tool for interpreting omics data. The Innovation, 2(3), 100141. [doi:10.1016/j.xinn.2021.100141](https://doi.org/10.1016/j.xinn.2021.100141).

Yu G, Wang L, Han Y, He Q (2012). clusterProfiler: an R package for comparing biological themes among gene clusters. OMICS: A Journal of Integrative Biology, 16(5), 284-287.[doi:10.1089/omi.2011.0118](https://doi.org/10.1089/omi.2011.0118).

Yu G (2025). enrichplot: Visualization of Functional Enrichment Result. R package version 1.26.6, <https://yulab-smu.top/biomedical-knowledge-mining-book/>.

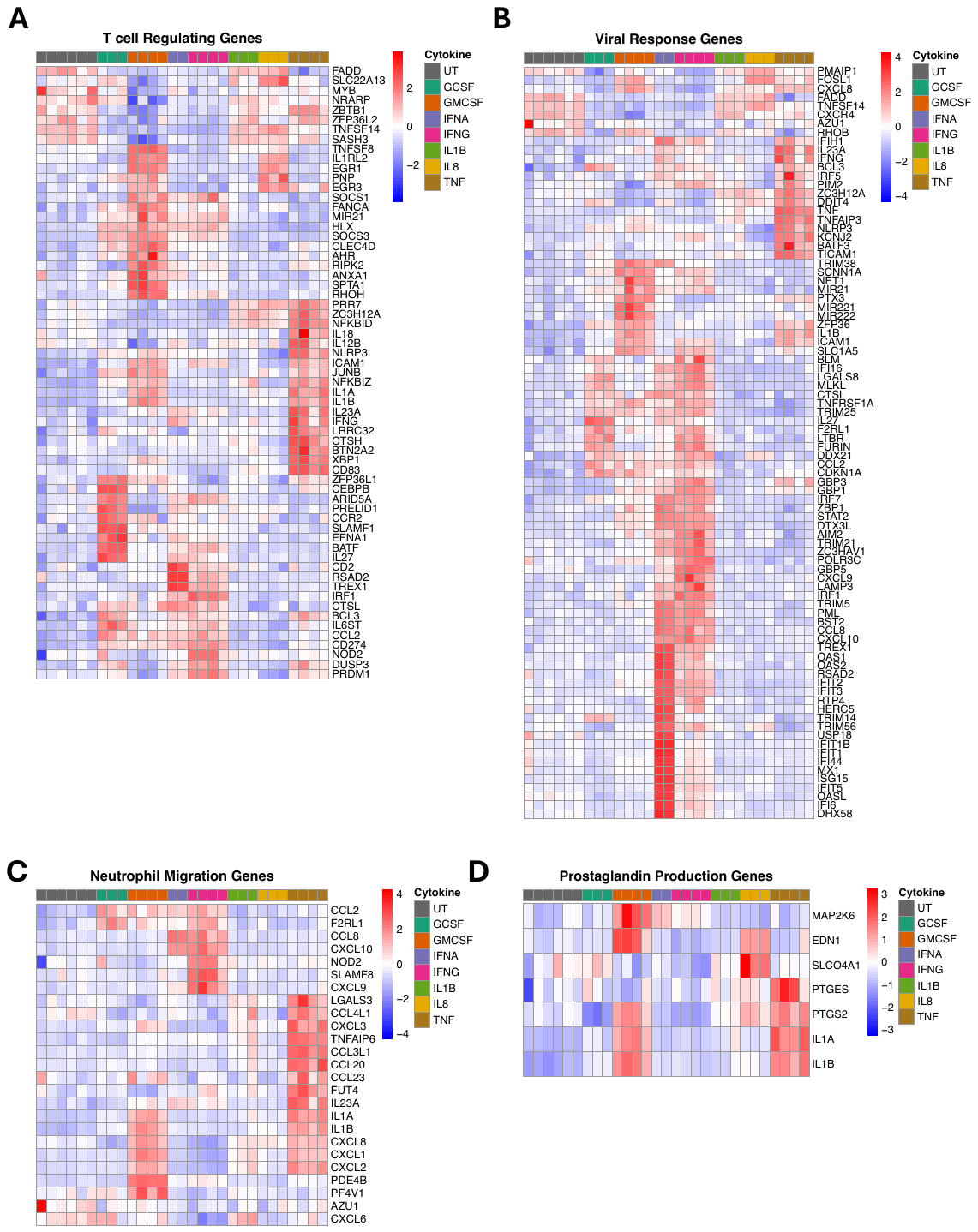

**Supplementary Figure 1. Heatmaps of genes common to GO BP terms**. (A) Genes enriched in ‘T cell regulation’, (B) Genes enriched in ‘Response to Virus’, (C) Genes enriched in ‘Neutrophil migration’, (D) Genes enriched in ‘Prostaglandin synthesis’.

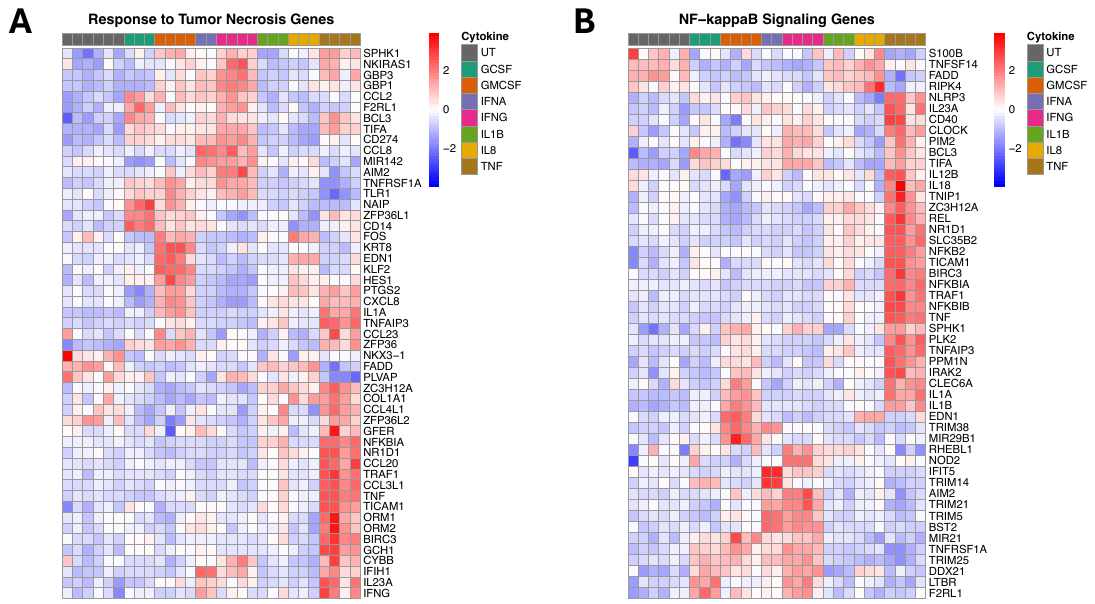

**Supplementary Figure 2. Heatmaps of genes common to GO BP terms**. (A) Genes enriched in ‘Response to tumor necrosis’, (B) Genes enriched in ‘NF-kappaB signaling’.
